# The Quantum Environment in Cryptochrome Enhances Light Absorption of FAD

**DOI:** 10.64898/2026.04.24.720615

**Authors:** Luc Wieners, Martin E. Garcia

## Abstract

Cryptochromes are blue-light photoreceptors that regulate circadian rhythms and, in some species, contribute to magnetic-field sensing through excitation of the flavin chromophore FAD. However, theoretical descriptions of cryptochrome photoexcitation usually treat only the chromophore quantum mechanically, neglecting quantum effects of the surrounding protein because of the computational cost of electronic-structure calculations. Here we present a computational framework that extends the quantum-mechanical region from the chromophore alone to approximately 1,200 atoms, enabling explicit treatment of the surrounding amino acids and cofactors. The approach combines relaxed-accuracy real-time time-dependent Hartree–Fock simulations with a post-processing correction that compensates for systematic errors.

The framework resolves the spatial origin, wavelength and oscillator strength of individual absorption processes throughout the quantum region. Our calculations reveal that the quantum-mechanical protein environment substantially enhances the light absorption of FAD, demonstrating that neighboring residues actively modulate the photophysical response of cryptochrome rather than merely providing a structural scaffold.

## Introduction

Cryptochromes are proteins which play a main role in sensing light in many plants and animals. This is achieved by their chromophore molecule FAD which absorbs blue to ultraviolet light. Light absorption in this ligand leads to a cycle of chemical reactions which causes conformational changes in the protein. The exact changes in the protein and how it signals incoming light to the organism are still an area with ongoing research [1–6].

For these reasons, cryptochrome was identified as an important protein in maintaining circadian rhythms in organisms [7]. Next to acting as a photoreceptor it also plays a role in inhibiting the transcription of DNA. Furthermore, cryptochrome is involved in sensing magnetic fields in many species [8–10]. The sensing is proposed to happen via a radical pair mechanism [11] between FAD and a neighboring residue which is a phenomenon only observed in cryptochrome.

Consequently, cryptochrome and its ligand FAD has been a target for experimental [12, 13] and theoretical [14–17] studies to gain better understanding of the underlying mechanism of light capture in this protein family. For the theoretical studies, a major challenge is the size of the protein cryptochrome as it consists of around 8,000 atoms, leaving the complete protein out of reach for a fully quantum-mechanical calculation of its absorption. Focusing on the ligand FAD is therefore sensible as it has the central role in the photocycle process. However, the molecule FAD can already be considered quite large for spectroscopic studies since it contains around 85 atoms, depending on the oxidation state. While one study reports a completely quantum-mechanical treatment of FAD [14], another approach is to investigate only the flavine group consisting of three aromatic rings with quantum mechanics [15, 16] as the absorption of ultraviolet and blue light is known to happen there.

While a reduction of the quantum region is excellent for increasing computational speed or to even make a study computationally feasible, it neglects the environment of the quantum region. Therefore, this approach is commonly coupled with another way of accounting for the environment or the rest of the molecule. A common method is to combine molecular mechanics (MM) [18–21] and quantum mechanics (QM) [22, 23] which is called QM/MM [24].

Simulations of this kind provide the ability to study very large systems and are therefore considered as state-of-the-art in large-scale spectroscopy. The drawback of the QM/MM methodology is that the MM part does not describe the system at a subatomic level and therefore brings limitations. Consequently, a fully quantum-mechanical framework is needed to study environment effects with the inclusion of electronic degrees of freedom.

Here, we achieve this by using the time-dependent Hartree-Fock method (TDHF) with a minimal basis set which allows fast calculations even for very large systems [25]. This method is combined with a post-processing to compensate systematic errors which occur due to this approach; further details about the computational method are provided in the methods section. As an additional point next to presenting calculations with a large-scale quantum region, we also investigated the differences between cryptochrome absorption among species by performing calculations for seven cryptochrome samples for which experimental reference spectra are available. Furthermore, the fine structure of absorption peaks is investigated by computing absorption spectra for structural data of FAD molecules obtained from the protein data bank (PDB) [26].

Finally, we present three examples for absorption spectra calculations of the chromophore FAD in combination with another light-absorbing system, studied in the same system with a quantum-mechanical environment. These three systems are a chain of tryptophan residues involved in the electron transfer process in two different species and the light-collecting cofactor MTHF which supports the light coupling in some cryptochromes. In these systems, 1,100 to 1,300 atoms are treated quantum-mechanically which makes them – to our knowledge – the largest quantum region used for studying light-harvesting proteins.

## Results

### Environment creation

The first two goals in this study are the investigation of the effects of including environment in the quantum-mechanical region and of studying the difference in absorption between species compared to experimental results. A starting point for all calculations is always the chromophore in the form of the molecule FAD. For investigating environment-based effects, we consider different configurations of the environment around the ligand: one configuration with no environment, one with an environment of three and one with five Angstrom (see Figure 1). The procedure of calculating these environments is described in the supplementary information.

**Figure 1:**
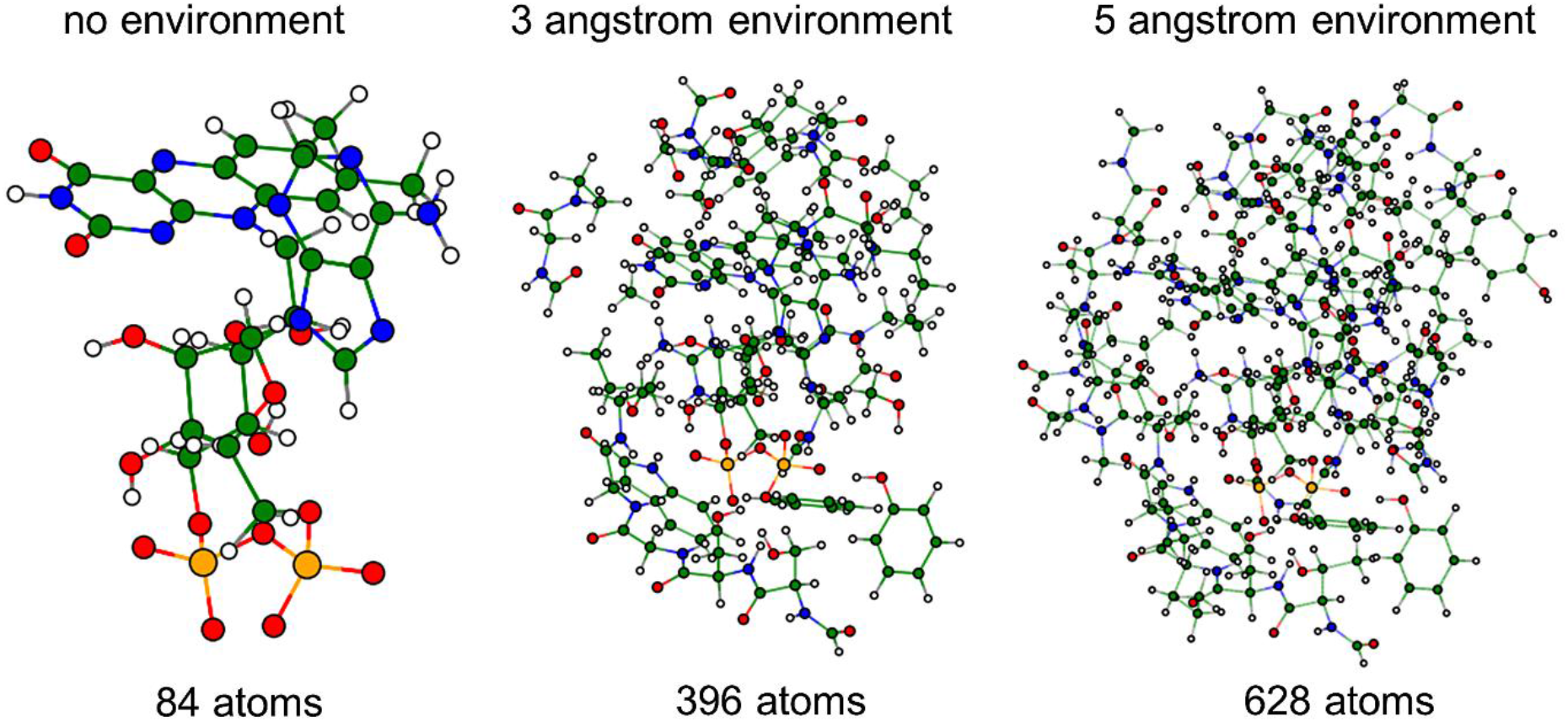
Atomic environments of FAD in cryptochrome. The different environments used for computing the absorption spectrum of cryptochrome are shown: no environment, three and five Angstrom environment. The corresponding atomic configurations, which include additional atoms to ensure sensible boundaries, are depicted for one of the seven systems (PDB 9LPG [27, 28]).

### Computational and experimental absorption spectra

First, we want to compare the computed spectra of seven cryptochrome structure samples from different species (the selection process is described in the methods section) with their respective experimental spectra and then investigate the effects of different environment sizes. For a comparison with experimental spectra, it is common practice to shift the theoretical spectra by a constant wavelength to account for systematic errors [14, 16, 29, 30]. Here, this calibration is done with the spectra obtained using the largest environment as this calculation should represent the biological system the best. A shift of 140 nm was found to be optimal. Note that we expect high shifts due to the use of the Hartree-Fock methods which systematically overestimates HOMO-LUMO gaps leading to absorption at lower wavelengths. In Supplementary Table 1, the positions of both peaks are given for the experimental and theoretical spectra with the discussed shift used for the latter. Here, an attenuation factor of 0.005 was used in the Fourier transform for a better resolution of the peaks.

If a smaller attenuation factor is used in the Fourier transform, we find that the major absorption peak of FAD at around 370 nm consists of two peaks in the theoretical spectrum. These are however not related to the two peaks in the experimental spectrum. Instead, one of the absorption peaks is caused by the adenine at the adenosine diphosphate (ADP) end of the FAD molecule. In the calculations, these two peaks overlap since the adenine, which absorbs at a lower wavelength than the flavine, would need a different shift as the error of its absorption peak is smaller than for the flavine group.

This stresses the importance of reference calculations when using the RT-TDHF framework to adequately interpret absorption peaks. Consequently, one has to keep in mind that the first absorption peak has a strength which is about twice as high as expected due to the overlapping absorption of the adenine. If this is accounted for, both relevant peaks have a similar intensity as it is the case for the experimental spectra.

Next, we investigate the difference in the absorption spectra for different environment sizes. This is shown in Figure 2 where the experimental and theoretical spectra are presented for all seven samples for the cases of no environment, 3 Angstrom and 5 Angstrom environments. It can be observed that the peak positions of the computed spectra agree well with the experimental peak positions in all environment configurations and not just for the 5 Angstrom environment which was used for the calibration. Consequently, for the case of FAD embedded in cryptochrome, the environment appears to not influence the positions of the absorption peaks of the chromophore in a significant way. This result is in good agreement with experimental data since the experimental cryptochrome absorption spectra used in this study agree strongly with the spectrum of isolated FAD [31].

**Figure 2:**
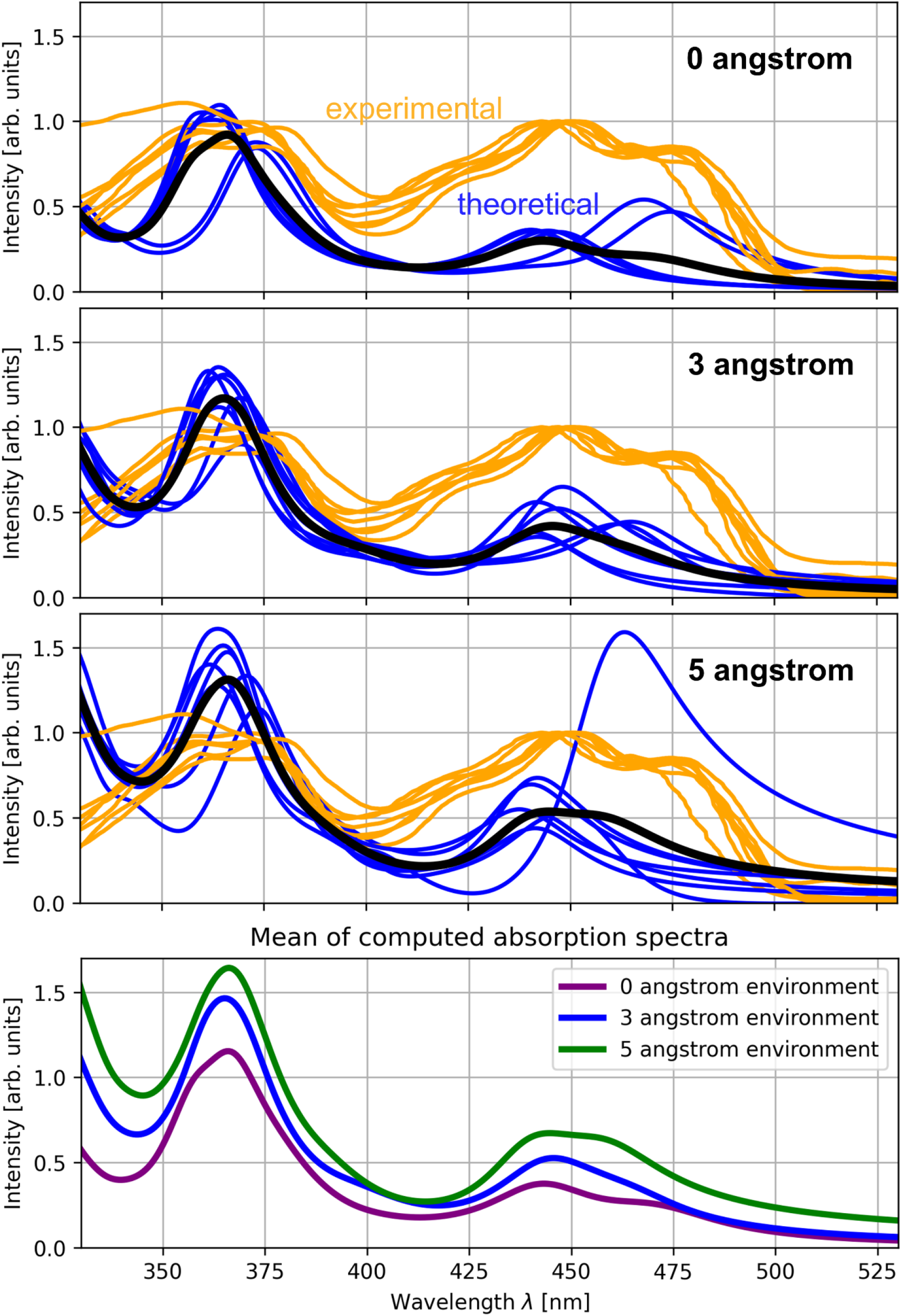
Absorption spectra in different environments. Comparison between experimental and theoretically obtained absorption spectra. The experimental spectra from the seven studied samples are shown in orange and the theoretical samples in blue, with the mean of the theoretical samples visualized as the black line. The top, middle and bottom panel show a comparison for environment sizes of 0, 3 and 5 Angstrom, respectively.

In addition to the absorption peak positions, the intensity of the absorption can be investigated over different environment sizes. In Figure 2 the theoretical spectra (in blue) show an increase in intensity for larger environment sizes, both for the step up from 0 to 3 Angstrom and from 3 to 5 Angstrom. One sample shows a very strong absorption at the second absorption peak of FAD for the 5 Angstrom environment. The intensity differences are shown in more detail in Supplementary Figure 1 where the three environment configurations are visualized in comparison with the respective experimental spectrum for each of the samples individually. An increase of the intensity of the absorption between 0 and 5 Angstrom environments can be seen for all systems at both major absorption peaks. In the bottom right panel of Supplementary Figure 1, all absorption curves are visualized together to illustrate this trend.

### Details of the FAD absorption spectrum

Another observation relevant for studying the absorption profile of FAD in detail is the fine structure of the absorption peaks. The two major absorption peaks of FAD are observed around 370 nm and 450 nm. However, the first peak is a double peak with absorption maxima at 355 nm and 375 nm, with the absorption at 375 nm being slightly higher than the absorption at 355 nm. Furthermore, the second peak consists of absorption at three different wavelengths. It has its overall maximum at 445 nm and a second maximum at 475 nm as well as a weaker peak at 425 nm.

These fine structure features are however not visible in computational studies – for both structures (the double and the triple peak) always only one maximum is reported (note that one peak of the double peaks in this study is caused by the ADP end of FAD). Therefore, we performed calculations for a larger set of existing structures of FAD molecules bound into cryptochrome. This list and its PDB accession codes are discussed in the methods section. In total it contains 28 structures for which the light absorption was computed in the same way as for the list of seven samples discussed above. In these calculations, no environment was used to focus on the absorption spectrum of FAD itself.

The results of these calculations are depicted in Figure 3 where all individual spectra and a mean spectrum of the FAD molecules is shown and compared to experimental spectra. Here, we observe a grouping of the absorption peaks at two different wavelengths for both of the two major absorption peaks of FAD. This results in both of the absorption peaks being split into two when the mean of all absorption spectra is computed.

**Figure 3:**
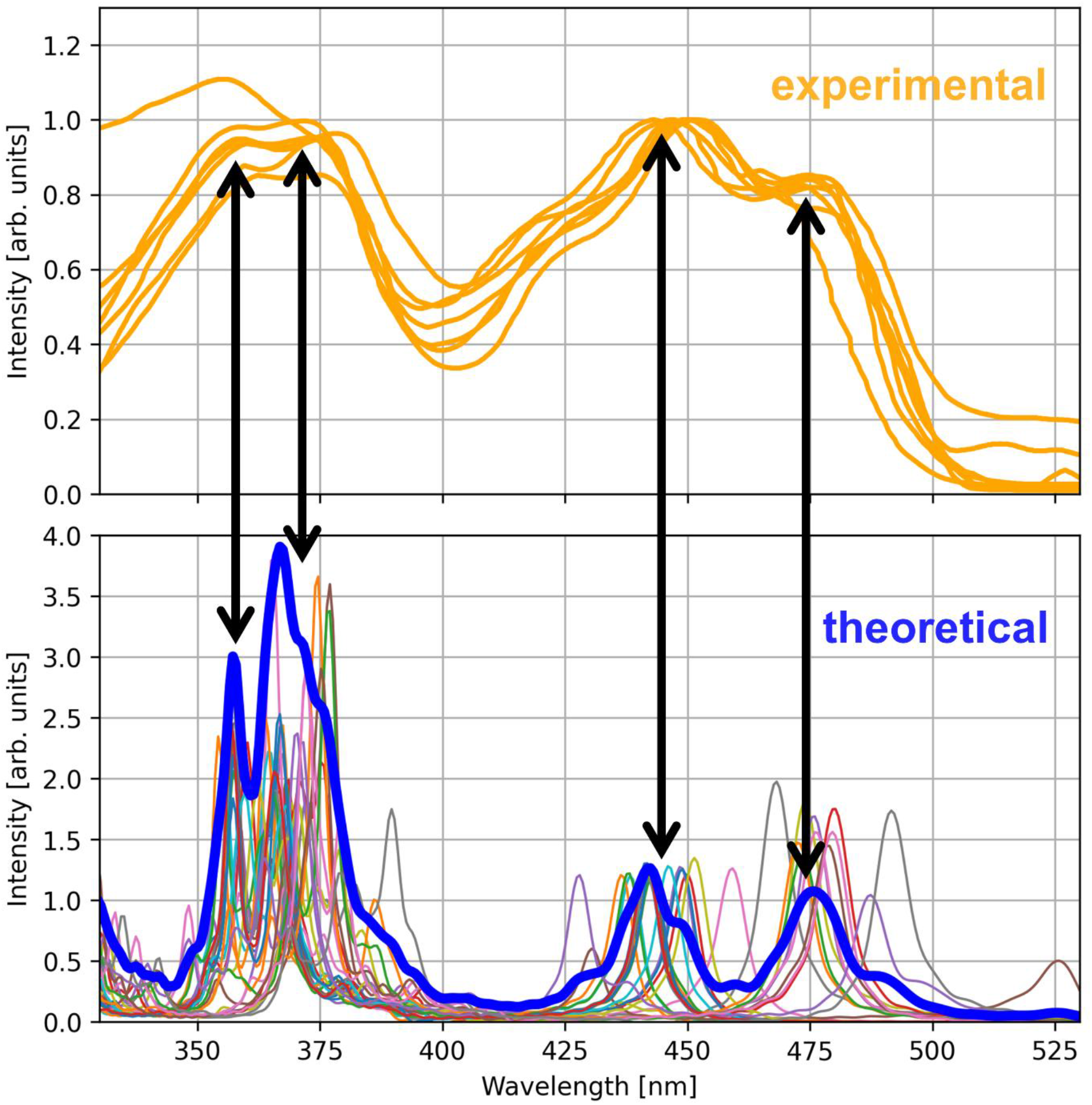
Details of the FAD absorption spectrum. Investigation of 28 FAD molecule structures bound to cryptochrome from the Protein Data Bank (PDB) [26] and comparison with experimental spectra. Top: Seven experimental absorption spectra from FAD in cryptochrome (see also Figure 2). Bottom: The individual spectra are shown as colored lines and the average is shown as a blue line. For visualization purposes, the average is multiplied by a factor of four. The black arrows connect the absorption peaks from the calculation with the peaks from the experiment. For a sharper visualization of the absorption peaks, an attenuation factor of 0.003 was used in the Fourier transform for these calculations.

Especially for the absorption peak at 450 nm, the splitting can be observed clearly in Figure 3 as well as in Figure 2 (most prominently in the top panel) and in Supplementary Figure 1 in the bottom right panel. This leads to the conclusion that different FAD structures lead to slightly different absorption peaks which has to be caused by the conformation of the FAD molecule since no environment is considered in the calculations presented in Figure 3.

### Systems with multiple absorbing molecules

Finally, we want to test the performance of the presented framework for larger systems in cryptochrome, exceeding system sizes of 1,000 atoms. For this purpose, we chose three systems where additional structures near the chromophore molecule FAD got investigated. First, we studied four tryptophan residues (the so-called “tryptophan tetrad”) [32–34] which are relevant for the electron transport pathway mechanism in the FAD photocycle and also show absorption at lower wavelengths. This was done for two species: *Drosophila melanogaster* (fruit fly) as a model organism and *Columba livia* (rock pigeon) as an organism that utilizes FAD and the tryptophan tetrad for magnetoreception. The PDB accession codes are 4GU5 [35, 36] and 6PU0 [37, 38], respectively.

Second, we investigated a cryptochrome with two ligands, FAD and MTHF from *Arabidopsis thaliana* (thale cress), PDB 2J4D [39, 40]. Here, MTHF is considered to serve as a light-collecting ligand to support the photocycle of FAD. The three considered systems have sizes of 1227, 1291 and 1145 atoms, respectively.

For the tryptophan residues it has to be considered that the shift of the absorption profile is different from the scaling factor in FAD. Therefore, the shift for tryptophan was computed individually by calculating the absorption spectrum of a single tryptophan molecule. This results in a shift of 50 nm. Due to the shift of FAD being 140 nm, the absorption maxima of FAD (375 nm) and tryptophan (280 nm) can be found close to each other, similar to the absorption maxima of the flavine group and the adenine coinciding in the absorption spectrum of FAD.

To visualize where the absorption takes places, one can compute the Fourier transform not only for the total dipole molecule of the system but also for the dipole moments of the individual atoms [41]. This has been done for the three systems described above and allows to investigate where and at which strength the absorption takes place. In Figures 4 and 5 as well as Supplementary Figure 2 this is shown for the three studied molecules.

**Figure 4:**
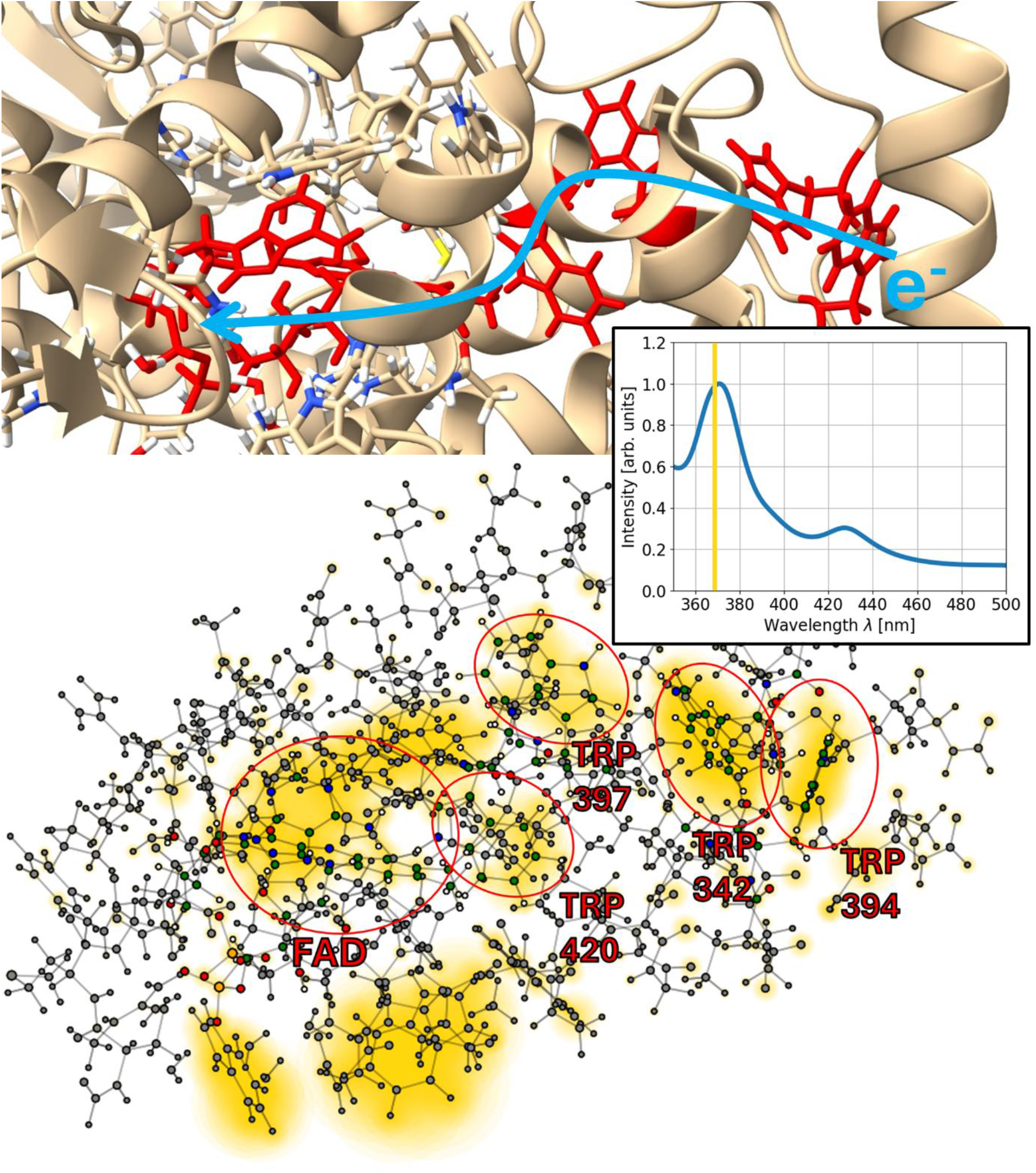
Absorption of the tryptophan tetrad in *Drosophila melanogaster* (fruit fly). Top: Visualization of the investigated system and its environment using ChimeraX [42–44]. The chromophore FAD and the four relevant tryptophan residues (W342, W394, W397, W420) involved in the electron transport are colored in red with the electron transport being indicated by the blue arrow. Bottom: The investigated system at an atomistic resolution. The core molecule and residues are marked in red with their atoms shown in color (all other atoms in grey). The absolute values of the Fourier transform of the dipole moments of all atoms are shown as yellow circles with the circle size corresponding to the respective value. They are displayed for a wavelength of 368 nm and indicate regions in which absorption may happen. The computed absorption spectrum is shown as an inset.

**Figure 5:**
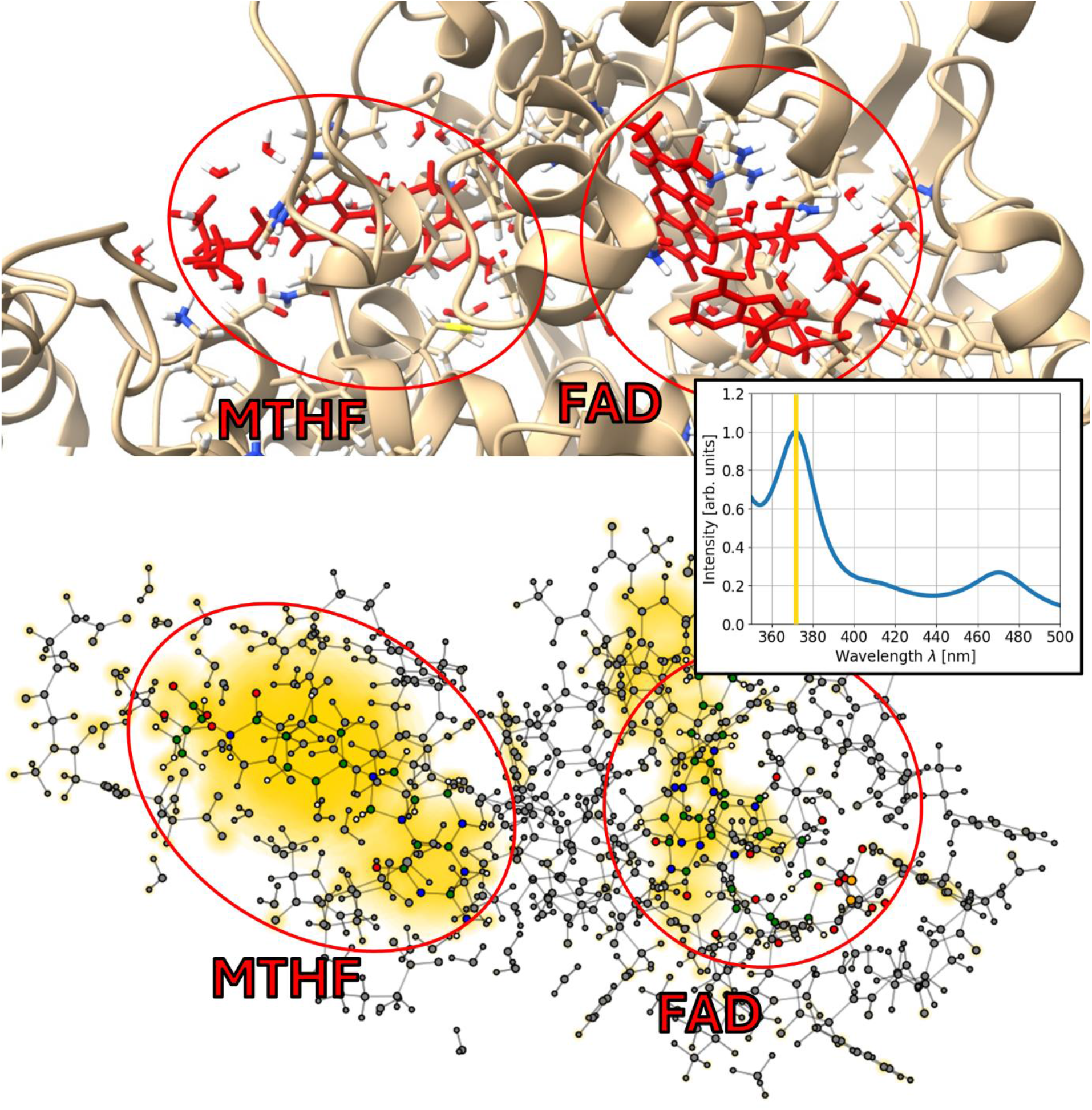
Absorption of MTHF and FAD. The absorption of the cofactor MTHF in cryptochrome is investigated in combination with the absorption of the chromophore FAD. Top: Visualization of the structure using ChimeraX [42–44] with the two ligands highlighted and circled in red. Bottom: Absorption at 372 nm is displayed in the same way as in the previous two figures. The computed absorption spectrum is shown as an inset.

In total, we observe absorption in FAD as well as in the tryptophan residues and MTHF. This is expected, since tryptophan is the amino acid with the strongest absorption and MTHF is known to absorb at around 370 nm. [39, 45, 46]

Consequently, the highest absorption peak in Figure 5 is attributed to FAD and MTHF, where MTHF shows stronger absorption than FAD, according to the visualization in Figure 5. For the systems with FAD and the tryptophan tetrad, the highest absorption is also found at 370 nm, this time as a combination of the absorption of FAD and the tryptophan residues (see Figure 4 and Supplementary Figure 2). Due to the absorption shift of tryptophan being 90 nm smaller than the shift of FAD, the absorption maximum corresponds to 280 nm for the tryptophan residues which is in agreement with the absorption peak position of isolated tryptophan.

For the systems with the tryptophan triad, the high absorption could be relevant for the electron transport as well since high absorption also corresponds to high polarizabilities.

The MTHF-FAD system shows a stronger absorption for MTHF than for FAD which corresponds well to the role of MTHF as the light-harvesting cofactor. It also matches experimental data of cryptochrome absorption spectra with FAD and MTHF where the absorption peak of MTHF dominates [39, 45].

## Discussion

The presented framework for quantum-mechanical absorption spectra calculations allows the study of biomolecules bound into proteins with a fully quantum-mechanical treatment of the environment in close proximity.

Certainly, a limitation of the described methodology is the higher error associated with the RT-TDHF calculations in a minimal basis. Therefore, the results often cannot be used without further investigation regarding the origins of the absorption peaks. Despite having multiple absorption overlaps at around 370 nm in this study, we found that using experimental reference spectra and reference calculations of isolated molecules, these errors can be controlled reliably. The presented protocol is designed to allow further studies of light absorption processes with a quantum-mechanical environment in many more systems since surroundings with a high polarizability in general have the potential to significantly influence the absorption spectrum. This has also been demonstrated by Frederiksen et al. who found that a polarizable embedding around the ligand FAD is necessary to perform high-accuracy spectral calculations in cryptochrome [15].

The option of an explicit quantum-mechanical treatment of environments is therefore promising to be of interest for a wide variety of absorption spectra calculations. This is especially the case if the surroundings are known to play an important role for the absorption spectrum or if parts of the environment are absorbing as well at a different wavelength. Furthermore, the presented method can be used to study the effect of solvents for larger structures with the QM region including the solvent.

In addition to this, larger quantum regions allow the treatment of multiple absorbing molecules and their environment in a single system, as demonstrated for MTHF and a chain of tryptophan residues in this study. A first step into studying the tryptophan residues in the electron transport chain could also be interesting for investigating magnetoreception since the chain of four tryptophan residues is postulated to play an important role in this topic [47, 48].

To conclude, we presented calculations of the light-sensing protein cryptochrome and its cofactor FAD while treating the close-range environment around the ligand fully quantum-mechanically. This presented method of QM environments for absorption spectra calculations was used to study seven structures of FAD bound to cryptochrome from multiple plants and animals. We observed that the region around the chromophore in cryptochrome leads to an amplification of the light absorption which in the protein is higher than in isolated FAD. Furthermore, we studied systems with multiple light-absorbing molecules, exceeding 1,000 atoms in total.

By doing so, we provided large-scale and systematic investigations regarding the light absorption of the chromophore FAD bound to cryptochrome regarding different environments, different species and in the presence of other light-absorbing molecules. Additionally, we provide a framework for studying absorption spectra of molecules with a quantum-mechanical treatment of the surrounding protein.

## Methods

### Experimental data for cryptochrome in animals and plants

For the procedure described above, the starting point is a structure file containing coordinates of the atoms in cryptochrome. In this part of the study, we focus on a comparison with preexisting experimental results from different groups [27, 28, 37, 38, 49–57]. For this purpose, we chose structures where an experimentally obtained absorption spectrum is available. The allocation of these configurations was done in the following ways. First, a search through all cryptochrome structures deposited on the RCSB protein data bank (PDB) [26] was performed. Then, those ones with FAD bound to cryptochrome and an experimental spectrum of the same molecule published in the paper corresponding to the PDB entry were selected. Additionally, samples with further functional complexes which changed the experimental absorption spectrum were avoided as well. In total, this leads to seven cryptochrome structure files with suitable experimental reference data for this study. The corresponding PDB entries are 9LPG [27, 28], 6K8I [49, 50], 6K8K [49, 51], 6X24 [52, 53], 6PU0 [37, 38], 6WTB [54, 55] and 8P4X [56, 57] and the respective species names are *Zea Mays*, *Arabidopsis thaliana*, *Arabidopsis thaliana*, *Arabidopsis thaliana*, *Columba livia*, *Drosophila melanogaster* and *Platynereis dumerilii*. This means that the dataset contains cryptochromes from maize, thale cress, rock pigeon, fruit fly and clam worm and consequently consist of both plants and animals.

### Computational method

In the absorption spectra calculations, we use the real-time time-dependent Hartree-Fock (RT-TDHF) method. This choice is motivated by a number of reasons. First, real-time methods scale at a computational cost of the third power of the system size while linear-response methods can scale up to the sixth power. Secondly, a central part in the computational complexity of realtime time-dependent Hartree-Fock or density functional theory is the construction of the Hamiltonian which needs to be done at every time step during the time propagation. For systems with a few hundred atoms this step is commonly much more time-consuming than the eigen-value calculation and therefore is considered a major bottleneck in computation time since the Hamiltonian matrix is computed thousands of times during a RT-TDHF or RT-TDDFT calculation [58]. The Hartree-Fock method allows a fast calculation of the Hamiltonian, especially the electron-electron interaction, since the integrals can be precomputed and stored which allows a fast assembly of the electron-electron tensor at each time step. This enables calculations with over 1,000 atoms.

A final advantage of RT-TDHF is that the Hartree-Fock method scales favorable for biophysical systems which have a lower density than for example systems in solid state physics. This leads to less integrals in total and a fast computation of the Coulomb and exchange interaction in the aforementioned building of the Hamiltonian. Our code NIMBLE [25] is focused on fast Har-tree-Fock calculations and filters out insignificant density interactions for further speedup. A detailed description of the used algorithms is given in the corresponding reference.

It should be discussed that this approach loses accuracy due the neglection of electronic correlation inherent to the Hartree-Fock method. Furthermore, we use the minimal basis (STO-3G) to maximize computational speed which introduces further errors. However, especially for the case of theoretical spectroscopy, a large part of these inaccuracies can be compensated by systematically comparison with reference data and reference calculations [29, 30]. The approaches to recognize, correct and control errors in this formalism are described in the following sections. These considerations come at the advantage of the mentioned ability to study larger systems previously out of reach as well as reasonable run times, making calculations accessible to smaller computation clusters or even personal computers, therefore saving time and energy.

For the quantum-mechanical time propagation, a protocol similar to [58] is used, utilizing the predictor-corrector time integration method. The propagation is done over 10,000 steps in total with a time step of 0.25 atomic units of time which leads to a total propagation time of 60.5 fs. The relatively long computation time ensures an accurate capture of dynamics at larger wavelengths and also allows a detailed resolution of the absorption peaks needed for quantitative comparisons with experiments. The Fourier transform uses an attenuation factor of 0.015 atomic units unless specified otherwise.

## Resource availability

Requests for further information and resources should be directed to and will be fulfilled by the lead contact, Luc Wieners. Coordinate input file data and result file data produced by the RTHF calculations and all original code have been deposited at Zenodo at 10.5281/zenodo.22942173 and are available from the lead contact upon request.

## Acknowledgments

L.W. and M.E.G. disclose funding and support for the research of this work from the German Research Association through the RTG 2749/1: “Biological Clocks on Multiple Time Scales”. Both authors would also like to acknowledge networking support by the COST Action CA21169, DYNALIFE, supported by COST (European Cooperation in Science and Technology).

Calculations were performed on the Linux Cluster of the University of Kassel and a cluster of the research group. L. W. thanks Barış Can Ülkü and Huleg Zolmon for useful discussions. Biomolecular graphics were performed with UCSF ChimeraX, developed by the Resource for Biocomputing, Visualization, and Informatics at the University of California, San Francisco, with support from National Institutes of Health R01-GM129325 and the Office of Cyber Infrastructure and Computational Biology, National Institute of Allergy and Infectious Diseases.

## Author contributions

Luc Wieners: Conceptualization, Data curation, Formal analysis, Investigation, Methodology, Project administration, Software, Validation, Visualization, Writing – original draft, Writing – review & editing Martin E. Garcia: Conceptualization, Funding acquisition, Investigation, Project administration, Resources, Supervision, Validation, Writing – review & editing All authors read and approved the final manuscript.

## Supplementary information

### General programme information

All absorption spectra calculations were performed using the real-time Hartree-Fock method implemented in our software NIMBLE. For the code and its detailed parametrization, we refer the interested reader to reference [25]. The STO-3G basis [59] is used and density-density interactions are truncated at a threshold of 10^−6^. Coulomb interactions are not truncated in the calculations.

Matrix operations are done with pytorch [60] while the computation of the electronic exchange matrix is done with a custom python routine which is just-in-time compiled and parallelized using the library numba [61], plots were done with the matplotlib library [62]. The RTHF calculations can also be parallelized among 3 compute nodes which calculate the responses for the electric field in x-, yand z-direction individually.

Most calculations were performed on compute nodes with 48 cores and took between a few hours to three days. For larger structures, it is necessary to have random access memory (RAM) in the range of 100-200 Gigabytes to ensure that the precomputed electron repulsion integrals, of which there are up to 7 billion for the largest calculation, can be stored. The time evolution of the dipole moments of the complete structure as well as for the individual basis functions are stored after the calculation is completed.

Coordinates and elements as well as the charge of the structure were precomputed and then read in again during the RTHF calculations. The procedure for precomputing these inputs is described in the following paragraph.

### Preparation of atomic coordinates

For the investigation of the differences between the three environment types used in this study (no environment, 3 Angstrom and 5 Angstrom), it is necessary to prepare the corresponding set of atomic coordinates and elements to perform the RTHF calculation. This is done by first identifying all atoms which have a distance to any of the atoms in the ligand which is less or equal than the given threshold (i.e. 3 or 5 Angstrom). Since only including these atoms would cut bonds on a non-physical way, additional atoms have to be added. This is done in a way so that the cutting of bonds changes the properties of the system as little as possible. For this reason, only single bonds between two carbon atoms are cut. The bond character is determined by computing the distance between the two atoms. Since the coordinates in this study are always taken from PDB files, there is very little deviation in the bond distances of bonds with the same character which makes this a reliable method for bond character determination. If a bond is being cut, the atom outside the added environment is replaced with a hydrogen atom with the bond distance chosen as the distance of a carbon-hydrogen bond.

In addition to preparing the atomic coordinates for the calculations with a protein environment, it is also necessary to determine the charge of the system. This is done by testing different charges in the form of adding or subtracting electrons from the system. For each of these tests, a few density matrix propagation steps are performed in the RTHF formalism and the charge of the FAD molecule is computed using Mulliken population analysis. The total charge is then selected among these test runs based on numerical stability and the charge of FAD where the latter should be equal to 408 (or in practice between 407.5 and 408). Note that 408 is two more than the sum of the atomic numbers of all elements in FAD. This is necessary due to the two phosphate groups in FAD which contain a negative charge each.

Consequently, the first criterion for choosing the correct electronic charge was the computed charge of FAD. In cases where multiple total charges lead to a suitable charge of FAD, the numerically stable case was chosen which was unique for all samples. The numerical stability was evaluated by computing the trace of the square of the density matrix which should correspond to the number of electrons in the system (except for a prefactor due to double occupations of orbitals). This is done routinely during RTHF calculations in NIMBLE to ensure that the time propagation runs stable – instable states are then identified due to a strong drift in the trace. The drift in the trace of the square of the density matrix can also be used to investigate the numerical loss or gain of electrons during the time propagation. The largest observed value in this study was found for the system containing FAD and the tryptophan tetrad for *Drosophila melanogaster* with a 5 Angstrom environment; here a deviation of 2.8 · 10^-8^ electrons/electron was observed after the completed time propagation over 60 fs, a value well within the scope of other quantum-mechanical time propagations [63].

The charges used in the calculations in this work are listed below, they are given in elementary charges i.e. 2 corresponds to removing 2 electrons.

– FAD without environment: –2 for all samples.
– CRY-FAD with 3 Angstrom environment: –3, –2, –2, –2, 1, 0, –1, for the seven samples in the order in which they are introduced in the main text.
– CRY-FAD with 5 Angstrom environment: –4, –3, –3, –3, –1, 0, 0, for the seven samples in the order in which they are introduced in the main text.
– TRP tetrad *Drosophila melanogaster*: 1
– TRP tetrad *Columba livia*: 0
– FAD-MTHF system: –6

For the study of the absorption peak positions of FAD without environment, 5 samples (sample numbers 3, 7, 8, 13 and 27 according to the list in the following section) were only numerically stable for charges other than –2. These samples were therefore not used since the focus was to simulate FAD absorption spectra at standard conditions.

Especially for the large-scale systems, the dipole moments were found to have a slight shift after the initial electric field pulse. This was corrected by subtracting the mean of the last 9,000 time steps of the dipole moment evolution for each coordinate before taking the Fourier transform.

Since the atomic coordinates are obtained from PDB files, only heavy atoms have explicit coordinates. The hydrogen atoms have to be added afterwards which was done using ChimeraX [42–44]. The hydrogen arrangement of the flavine group computed by ChimeraX does not match the structure of FAD in its oxidized form which is the molecule investigated in this study. For this reason, one hydrogen atom was deleted from the structures from ChimeraX to achieve the intended state.

### List of files for the investigation of FAD absorption details

For the comparison with experimental data and to investigate the details in the absorption profile of FAD, this study uses structures from the PDB [26]. The structures were obtained by searching for structures with the UniProt molecule names “Cryptochrome-1”, “Cryptochrome-2”, “Cryptochrome2”, “Cryptochrome DASH”, “Cryptochrome DASH, chloroplastic/mitochondrial”, “Cryptochrome/photolyase family protein” which are all listed molecule names that contain the word “cryptochrome”. Out of the PDB entries listed with these molecule names, only those with FAD as a ligand were selected. In total, this leads to 33 structures (as of late 2025).

In the following, a list of these 33 structures [27, 28, 35–40, 49–57, 64–100] is provided: PDB accession code, species name and classification of the relevant molecule according to the PDB entry are given.

1. 6LZ3, *Zea Mays*, Cryptochrome2
2. 9LPG, *Zea Mays*, Cryptochrome2
3. 2IJG, *Arabidopsis thaliana*, Cryptochrome DASH, chloroplast/mitochondrial
4. 2J4D, *Arabidopsis thaliana*, Cryptochrome DASH
5. 2VTB, *Arabidopsis thaliana*, Cryptochrome DASH
6. 7YKN, *Vibrio cholerae*, Cryptochrome/photolyase family protein
7. 8A1H, *Vibrio cholerae*, 6-4 photolyase (FeS-BCP, CryPro)
8. 9HNK, *Caulobacter vibrioides*, Cryptochrome/photolyase family protein
9. 9HNL, *Caulobacter vibrioides*, Cryptochrome/photolyase family protein
10. 9HNM, *Caulobacter vibrioides*, Cryptochrome/photolyase family protein
11. 9HNN, *Caulobacter vibrioides*, Cryptochrome/photolyase family protein
12. 9HNO, *Caulobacter vibrioides*, Cryptochrome/photolyase family protein
13. 9Q8F, *Caulobacter vibrioides*, Cryptochrome/photolyase family protein
14. 1NP7, *Synechocystis sp. PCC 6803*, DNA photolyase
15. 4I6G, *Mus musculus*, Cryptochrome-2
16. 6K8I, *Arabidopsis thaliana*, Cryptochrome-2
17. 6K8K, *Arabidopsis thaliana*, Cryptochrome-2
18. 6M79, *Arabidopsis thaliana*, Cryptochrome-2
19. 6X24, *Arabidopsis thaliana*, Cryptochrome-2
20. 7X0X, *Arabidopsis*, Cryptochrome-2
21. 7X0Y, *Arabidopsis thaliana*, Cryptochrome-2
22. 1U3C, *Arabidopsis thaliana*, Cryptochrome 1 apoprotein
23. 1U3D, *Arabidopsis thaliana*, Cryptochrome 1 apoprotein
24. 4GU5, *Drosophila melanogaster*, Cryptochrome-1
25. 4JZY, *Drosophila melanogaster*, Cryptochrome-1
26. 4K03, *Drosophila melanogaster*, Cryptochrome-1
27. 6LZ7, *Zea mays*, Cryptochrome-1
28. 6PTZ, *Columba livia*, Cryptochrome-1
29. 6PU0, *Columba livia*, Cryptochrome-1
30. 6WTB, *Drosophila melanogaster*, Cryptochrome-1
31. 7UD0, *Drosophila melanogaster*, Cryptochrome-1
32. 8DD7, *Drosophila melanogaster/Homo sapiens*, Methylated-DNA–protein-cysteine methyltransferase, Cryptochrome-1 fusion
33. 8P4X, *Platynereis dumerilii*, Putative light-receptive cryptochrome (Fragment)

### Peak positions of computed and experimental spectra

**Supplementary Table 1:** Peak positions of experimental and theoretical FAD absorption spectra. A shift of 140 nm is employed for the theoretical spectra (computed in a 5 Angstrom environment) and absolute and relative errors under this shift are given with respect to the experimental values of the same sample. All values are given in nm, except for the relative errors.

| Sample | Peak 1<br>exp. | Peak 1<br>theo. | Peak 2<br>exp. | Peak 2<br>theo. | Abs. error<br>peak 1 | Abs. error<br>peak 2 | % error<br>peak 1 | % error<br>peak 2 |
| --- | --- | --- | --- | --- | --- | --- | --- | --- |
| 1 | 365 | 364 | 443 | 439 | 1 | 4 | 0.3 | 0.9 |
| 2 | 367 | 374 | 449 | 460 | 7 | 11 | 1.9 | 2.4 |
| 3 | 369 | 373 | 447 | 456 | 4 | 9 | 1.1 | 2.0 |
| 4 | 369 | 367 | 451 | 440 | 8 | 11 | 2.2 | 2.4 |
| 5 | 366 | 364 | 447 | 441 | 2 | 6 | 0.5 | 1.3 |
| 6 | 368 | 365 | 451 | 442 | 3 | 9 | 0.8 | 2.0 |
| 7 | 365 | 363 | 451 | 438 | 2 | 13 | 0.5 | 2.9 |
| average | 367 | 367 | 448 | 445 | 4 | 9 | 1.0 | 2.0 |

### Absorption spectra by sample

**Supplementary Figure 1:**
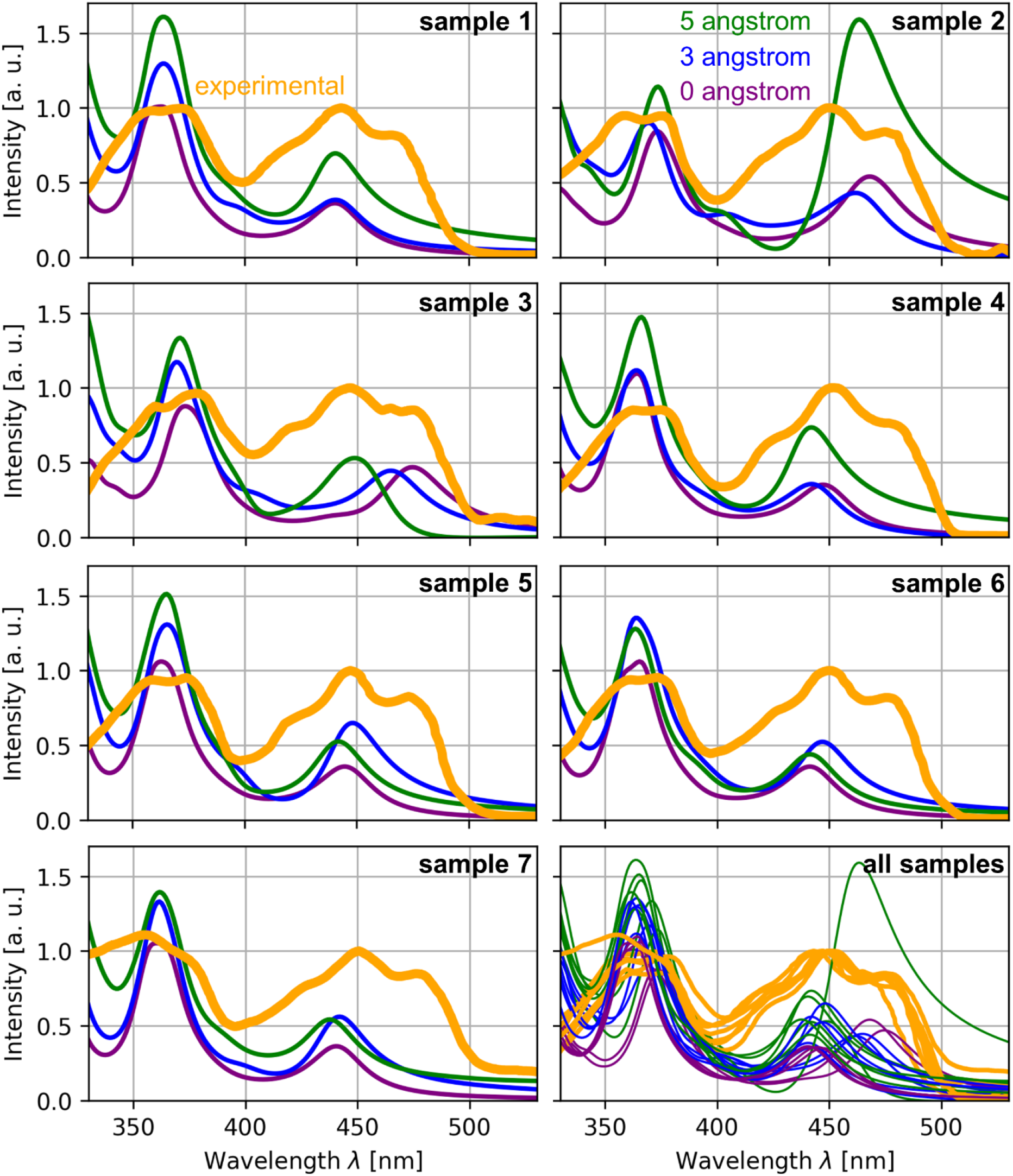
Absorption spectra for different cryptochrome samples. Comparison of spectra for different environment sizes and different samples. The spectra are shown for all 7 samples individually with the experimental and the three theoretical spectra displayed. In the bottom right panel, all spectra for all samples are visualized together.

### Tryptophan tetrad absorption in *Columbia livia*

**Supplementary Figure 2:**
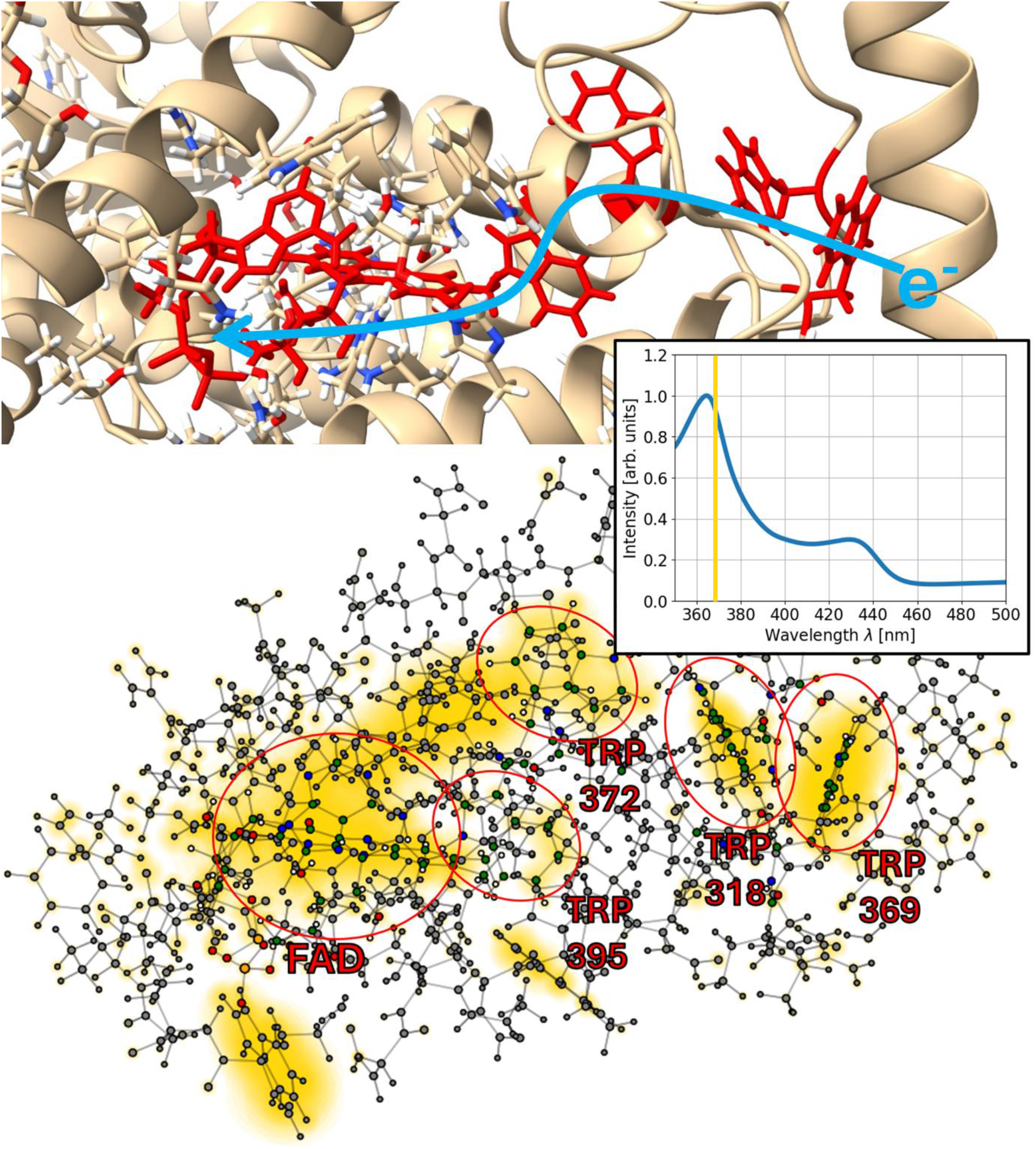
Absorption of the tryptophan tetrad in *Columbia livia* (rock pigeon). Top: Visualization of the investigated system and its environment using ChimeraX [42–44]. The chromophore FAD and the four relevant tryptophan residues (W318, W369, W372, W395) are highlighted. Bottom: A visualization of the absorption regions at 368 nm in the same way as in Figure 4. The computed absorption spectrum is shown as an inset.

